# TMT-based quantitative proteomic assessment of *Vicia sativa* induced neurotoxicity by β-cyano-L-alanine and γ-glutamyl-β-cyano-L-alanine in SH-SY5Y cells

**DOI:** 10.1101/2025.04.05.647336

**Authors:** Samuel Riley, Vy Nguyen, Rudrarup Bhattacharjee, Pei Qin Ng, Tarin Ritchie, Ian Fisk, Jozef Gecz, Raman Kumar, Iain R. Searle

## Abstract

β-Cyano-L-alanine (BCA) and γ-glutamyl-β-cyano-L-alanine (GBCA) are the primary antinutritional compounds present within the seeds of the high protein and drought tolerant orphan legume *Vicia sativa*. Evidence of neurotoxicity is limited to symptom analysis from animal feed trials whilst the molecular mechanisms underlying suspected neurotoxicity in monogastric animals are largely unknown. In this study, we first optimised an *in vitro* cell-based assay for rapid testing of BCA and GBCA toxicity in the retinoic acid differentiated SH-SY5Y human neuroblastoma cells. Using this system, we then performed proteomics analyses to determine dysregulated expression of proteins in BCA or GBCA treated of differentiated SH-SY5Y cells. Our findings indicate that BCA affects expression of proteins involved in DNA damage and translation whilst GBCA treatment causes dysregulation of those involved in mitosis and cell cycle. Following BCA treatment, we identified changes in many proteins previously suggested to have close association with neurodegenerative diseases including amyotrophic lateral sclerosis and Alzheimer’s disease as well as cancers. Following GBCA treatment, dysregulation of proteins involved in apoptosis pathways was observed. Finally, the lack of common dysregulated proteins and pathways in BCA or GBCA treated cells indicated that they most likely cause neurotoxicity via distinct mechanisms.

**Significance:** Antinutritional compounds or toxins limit the use of many potential grain crops including *V. sativa* (Common Vetch), and BCA and GBCA are the two principal toxic compounds in Common Vetch grain. Vetch grain toxicity is associated with neurotoxic symptoms in animals. Characterizing the proteome of BCA- and GBCA-treated neural cells gives insights into the excitotoxicity mechanism and identification of biomarkers for screening higher quality grain. Quantitative tandem mass tag (TMT) mass spectrometry-based proteome profiling of BCA treated neural cells characterized 6,827 proteins, of which 26 were significantly up-regulated and 73 significantly downregulated. However, TMT proteome analysis of GBCA treated cells identified 76 significantly up regulated and 86 downregulated proteins. These results provide insight into the toxicity of BCA and GBCA and enable future identification of putative direct protein targets. We describe for the first time BCA playing a role in modulating the expression DNA damage proteins and translation, and GBCA playing a role in modulating the expression of mitosis and cell cycle proteins, elucidating the mechanisms of plant-derived toxins in mammalian cell neurotoxicity

**Highlights:**

- Established *in vitro* cell-based toxicity assay for β-cyano-L-alanine and γ-glutamyl-β-cyano-L-alanine.
- β-Cyano-L-alanine treatment dysregulated many proteins associated with DNA damage in retinoic acid differentiated SH-SY5Y cells.
- γ-Glutamyl-β-cyano-L-alanine treatment dysregulated proteins associated with cell cycle.

## Introduction

The world population is rapidly increasing and is expected to reach nine billion by 2050 [1]. A population of this size will require a 60 % increased agricultural capacity [2] to supply both sufficient caloric energy and nutritious high-quality protein. Current global protein supply is reliant on the livestock industry, where the process of husbandry has low land use efficiency [3] and significant greenhouse gas emissions [4]. To negate these impacts, plant protein sources from chickpea (*Cicer arietinum*), faba bean (*Vicia faba*), lentil (*Lens culinaris*), pea (*Pisum sativum*) and soybean (*Glycine max*) are currently utilized and are proposed to partially fulfil the increased protein requirements. Such leguminous crops are highly valued in agriculture for their capacity to fix atmospheric nitrogen into bioavailable forms, reducing the need for chemical nitrogen inputs. Integrating legumes in crop rotations also limits soil erosion, suppresses weed growth and increases soil organic matter. Research has focused on increasing the adaption and hence increased yield of these legume species by utilising diverse germplasm to current agricultural areas and more recently, identifying new legume species adapted to surviving in marginal agricultural environments. One key species under investigation is *Vicia sativa* (Common Vetch) which shows great potential as a human food crop [5].

Common vetch is an under-utilised leguminous crop that has the potential to be a prominent global pulse in fringe agricultural systems [6]. *V. sativa* is recognized for its superior drought tolerance compared to other legumes [7], has high seed crude protein (24-32 %) [8] and low lipid content (1.5-2.7 %) [9]. Despite the recognised benefits of common vetch towards reducing soil erosion, suppressing weed growth and increasing soil organic matter when used as a manure, its adoption as a human food crop has been limited due to the presence of antinutritional factors in the seed, namely the dipeptide γ-glutamyl-β-cyano-L-alanine (GBCA, 2.6 %) and the free amino acid β-cyano-L-alanine (BCA, 2.6 %) [10]. Both compounds are acutely toxic to monogastric species [8] and upon addition in animal feeds cause symptoms highly associated with neurotoxicity including hyperactivity, convulsions, tremors, rigidity, prostration, and in some cases death, e.g., in rats and chicks [11–13]. The excitotoxicity mechanism through which BCA acts is not clearly understood. In animal trials, pigs, and rats experienced neurotoxicity symptoms when eating a *V. sativa* supplemented diet, however HPLC analysis of brain tissue indicated presence of relatively low concentrations of BCA [14]. Notably, analysis investigating if BCA acts through release of cyanide ions observed no inhibition of NADH dehydrogenase (complex I) or cytochrome or oxidase (complex IV) activity [15]. These results suggest the observed neurotoxicity may be indirectly caused by BCA inhibiting distinct metabolic pathways [16]. BCA has been shown to inhibit the action of cystathionine γ-lyase (CSE), an enzyme responsible for catalysing cystathionine into cysteine that is eventually metabolised into glutathione, which acts as a sink for excess cysteine [12,17]. Disruption of this pathway can cause accumulation of reactive oxygen species, free radicals, and heavy metals, leading to cell necrosis and, dependent on the locale, symptoms typical of neurotoxicity. In summary, the excitotoxicity mechanism through which BCA acts remains to be clearly elucidated.

Currently, the exact concentrations of *V. sativa* toxins detrimental to human health and the precise mechanisms of BCA and GCBA neurotoxicity are unknown. High profile commentaries such as ‘a mess of red pottage’ [18] detailing the illegal substitution of de-hulled red lentils with physiologically similar but toxic *V. sativa* cultivar blanche fleur, and the fears of potential health risks, spurred rapid development of HPLC and LC-MS/MS protocols that facilitated successful identification of adulterated lentil products [19,20]. While these assays provide quantitative data for GBCA and BCA concentrations in the seed, they give no indication about the concentrations that would affect a biological system. Expensive and time-consuming feed trials have determined the safe amount of *V. sativa* grain that could be used as a part of feed for specific monogastric species, for example, for pigs, layer hens, rabbits, and broiler chickens, relatively low levels (100-300 g/kg) [21], [22]. One option to animal trials is *in vitro* cell assays. *In vitro* cell assays are commonly used for drug discovery and metabolic studies as they generate significant biologically relevant information than that by analytical biochemical assays, and reduce the amount of animal tests required [23].

In this study, we investigated the effects of BCA and GCBA on *in vitro* cultured neuronal cells derived by differentiating the routinely used SH-SY5Y human neuroblastoma cell line. In addition, we investigated dysregulation of proteins underlying the specific toxicity mechanisms impacted in BCA or GBCA treated differentiated SH-SY5Y cells by assaying qualitative and quantitative proteome using tandem mass tag (TMT) approach. In addition, we developed an *in vitro* cell assay for rapid detection of BCA and GBCA from vetch seed extracts.

Our overall aims were to identify neuronal pathways that are affected by BCA and GBCA toxins and to explore the possibility of developing a high throughput *in vitro* cell assay for rapid testing of BCA and GBCA in vetch seed extracts. Deployment of the rapid assay allowed expeditiousdetection of a zero-toxin vetch accessions and facilitated the development of vetch as a new sustainable food source for human and animal.

## Materials and methods

Seeds of *V. sativa* (acc. Timok, Lov-2) were procured from S&W Seed Company (Longmont, CO, USA). 100×17 mm or 60×16 mm Nunc EasYDishes and Nunc Cell-Culture treated 12 well plates were purchased from ThermoFisher Scientific (Waltham, MA, USA). γ-Glutamyl-β-cyano-L-alanine was purified as described [24] and the purity was assessed by NMR. Diethyl ethoxymethylenemalonate (DEEMM), acetonitrile (HPLC grade), All-*trans* retinoic acid (RA), human brain derived neurotrophic factor (BDNF) and β-cyano-L-alanine were purchased from Sigma–Aldrich (St. Louis, MO, USA). Dulbecco’s Modified Eagle: Nutrient Mixture F-12 (HAM) 1 : 1 medium (DMEM/F-12), and fetal bovine serum (FBS) were obtained from Gibco Invitrogen (Waltham, MA, USA). Alamar Blue Cell Viability Reagent was purchased from ThermoFisher Scientific (Waltham, MA, USA). Penicillin-Streptomycin and GlutaMAX was from Gibco (Waltham, MA, USA).

### SH-SY5Y neuronal cell differentiation

SH-SY5Y cells (ATCC, Virginia, USA) were maintained in DMEM: F12 complete medium supplemented with 10 % heat inactivated fetal bovine serum (HI-FBS) and 1 % Penstrep. Cells were cultured in a humidified, 5 % CO_2_, 37 °C incubator.

SH-SY5Y were differentiated into neuronal cells as described [26]. In brief, SH-SY5Y cells were seeded in DMEM, 10 % HI-FBS and 1 % penstrep for 18 h before culturing in DMEM, 10 % HI-FBS and 0.1 % RA (10 mM) for five days, following three days in DMEM, 0.1 % BDNF. Successful differentiation of SH-SY5Y cells was confirmed by western blotting using differentiation marker neurofilament H (Figure 1).

**Figure 1.**
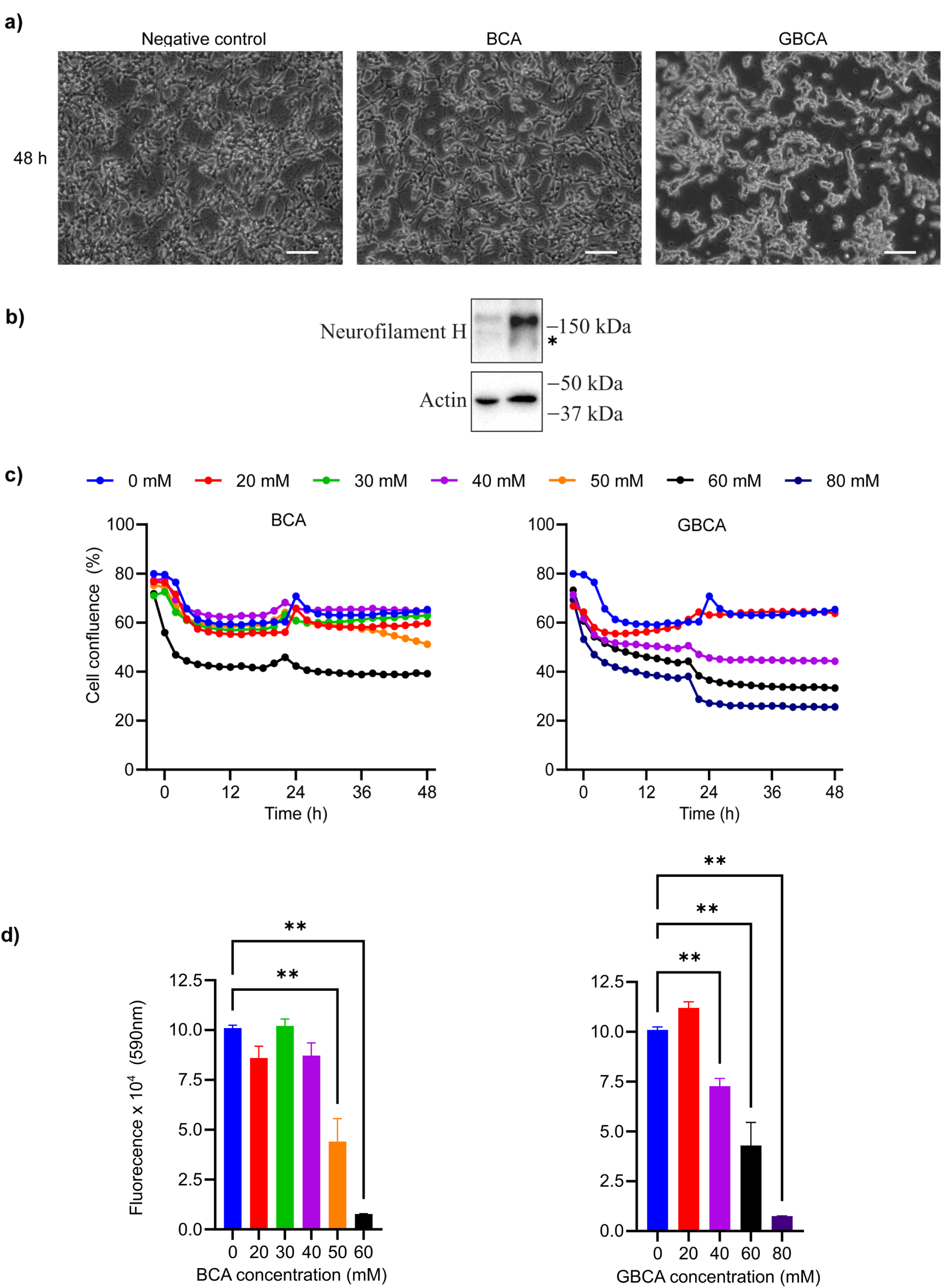
Cell viability of differentiated SH-SY5Y cells is affected by BCA or GBCA treatment. a) Cell morphology of control, BCA (50 mM) or GBCA (50 mM) treated differentiated SH-SY5Y cells after 48 h. Scale bar = 200 µm. b) Validation of SH-SY5Y neuronal differentiation using western blotting for the differentiation marker Neurofilament H. Actin was used as a housekeeping protein loading control. *, a non-specific or degradation product. c) Real-time cell viability (measured by cell confluence using IncuCyte S3 system) of differentiated SH-SY5Y cells treated with different concentrations of BCA or GBCA. d) Relative cell viability after 48 h of control, BCA or GBCA treated differentiated SH-SY5Y cells as determined by Alamar Blue assay. Data shown as mean ± SEM of three biological replicates; ** p<0.01.

### BCA and GBCA treatment of differentiated SH-SY5Y cells

SH-SY5Y cells were seeded at 5×10^3^ cells per well in a 96-well Nunc plate and were differentiated as described above. The culture media was then replaced with 100 μL of DMEM: F12 complete medium, 0.1 % BDNF, and GBCA or BCA at concentration from 20 to 80 mM. Cells were cultured in a humidified, 5 % CO_2_, 37 °C incubator for 48h before cell viability was measured.

### BCA and GBCA treatment for mouse primary neurons

Mouse primary neurons were isolated as described previously [25] and were seeded at 2×10^6^ cells in 1mL neurobasal A with 2 % B27 supplement, 1 % penstrep and 1 % glutamax (neural feed) per well, in a 12-well CoStar plate (Corning Inc, Corning, NY, USA). Cells were maintained with half media changes every three days. On the seventh day after seeding, cells were treated with BCA or GBCA (final concentration ranging from 20 mM up to 80 mM) in neural feed. Cells were cultured in a humidified, 5 % CO_2_, 37 °C incubator for 48 h before cell viability was measured.

### Cell proliferation assay

To determine the cell proliferation rate, differentiated SHSY-5Y cells plated in 96-well plates and mouse neurons plated in 12-well plates were loaded into the live-cell imaging system IncuCyte S3 (Sartorius AG, Göttingen, Germany) and an average of four images per well, at 10× magnification, were recorded every 2 h for 48 h. The data were used to quantify the cell surface area coverage (confluence values) using the IncuCyte Confluence software (version 1.5) (Sartorius AG, Göttingen, Germany).

### Cell viability determination

Cell viability was determined using Alamar Blue fluorescence assay (cat. DAL1025, Invitrogen). In brief, cells were incubated in 10 % Alamar Blue in DMEM: F12 complete medium in dark at 37°C for 12 h. Emitting fluorescence from live cells was measured at 590 nm to determine the cell viability. All data are average of biological triplicates.

### Assessment of *V. sativa* seed extract toxicity

#### V. sativa seed extract preparation

*V. sativa* seed extraction was performed using a modified method described previously [27].

*V. sativa* seed (Timok and Lov-2 cultivars) was grounded using a coffee grinder (Breville BCG200, Alexandria, Australia). The milled seed was filtered through 200 µm nylon-mesh and 200 mg of the milled seeds were mixed with 1 mL of 30 % ethanol at room temperature for 1h. The supernatant was separated from the mixture by centrifugation at 11,500 × g for 10 min and the extraction with 30 % ethanol was repeated for the pellet twice. A total of 3 mL supernatant was pooled and freeze-dried using Alpha 2-4 LD plus (Martin Christ, Osterode am Harz, Germany). For vetch seed extract that required filtering, seed extracts were placed into a 3 kDa molecular weight cut-off Amicon Ultra-0.5 Centrifugal Filter Unit (Millipore, Burtlington, MA, USA) and centrifuged at 11,500× g for 10 min before freeze-drying. Freeze-dried samples were stored at -80°C for future use.

#### Testing the effect of V. sativa seed extract on differentiated SH-SY5Y cells

Seed extract supplemented media was prepared by resuspending the freeze-dried seed extract to the original pooled volume (3 mL) using DMEM: F12 complete medium supplemented with 0.1 % BDNF. The solution was filter-sterilised using 0.45 µm filter centrifuge tubes (Corning, Costar, Spin-X, Corning, NY, USA). The effect of seed extracts on the differentiated SH-SY5Y cells was tested as follows: SH-SY5Y cells were seeded at 5×10^3^ cells per well in a 96-well Nunc plate and differentiated as described above. The cell culture media was then aspirated and replaced with 100 μL of the seed extract supplemented media. For a negative control, DMEM: F12 complete medium supplemented with 0.1 % BDNF was used. GBCA (0–40 mM) supplemented DMEM: F12 complete medium with 0.1 % BDNF was used in positive control samples. The cells were cultured in a humidified, 5 % CO_2_, 37 °C incubator for 48 h before cell viability was measured.

#### Quantification of GBCA and BCA in seed extracts using HPLC-MS

The HPLC-MS system used in this study comprised a 1200 Infinity LC System (Agilent Technologies, Santa Clara, CA, USA) coupled to a 6230 time-of-flight mass spectrometer (Agilent Technologies). Electrospray ionisation in positive ion mode was used and Agilent Mass Hunter Data Acquisition, Qualitative Analysis and Quantitative Analysis software was used for method development and data acquisition.

Sample analysis using the HPLC-MS system is as follows: Freeze-dried samples were resuspended to the original pooled volume (3 mL) using 1 M borate buffer pH 9.0. To 1 mL of the resuspended sample, 0.66 µL of DEEMM and 10uL 25 mM L-2-aminobutyric acid (internal standard) were added and mixed. Samples were then derivatised at 50°C for 50 min before centrifuged at 18,500× g for 30 s to pellet any large particle if present. For each derivatised sample, 900 µL was transferred to LCMS vials and the vials were loaded onto HPLC-MS sample loader. Samples (10 µL) were injected for separation with an InfinityLab Poroshell 120 EC-C18 column (2.1×150 mm, 4 µm, Agilent Technologies) protected by a 2.1 mm guard cartridge of the same material. Elution was carried out using 0.5 % formic acid in MQ water (solvent A) and 0.5 % formic acid in acetonitrile (solvent B) in a binary gradient; starting at 25 % acetonitrile, increasing to 60 % over 11 min, further increasing to 90 % from 11 to 13 min which was maintained for 3 min; the initial conditions were recovered in 2 min. Column temperature and eluent flow were set at 22°C and 0.3 mL/min, respectively.

Peak area of derivatised BCA m/z = 284.27) and GBCA (m/z = 413.38) and internal standard (m/z = 273.28) were identified using Agilent Mass Hunter Data Acquisition; Qualitative Analysis and Quantitative Analysis software. The BCA and GBCA peak area were normalised by peak area of internal standard (analyte/internal standard), before BCA and GBCA concentration in the samples were predicted using a standard curve.

Samples with known amount of BCA and GBCA (ranging from 15 to 500 μM) in 1 M borate buffer pH 9.0 were analysed by HPLC-MS and the resulted data were used to construct standard curves to predict the BCA and GBCA concentration in the seed extract samples.

### LC-MS/MS analysis for proteomics

#### Sample preparation for TMT analysis

SH-SY5Y cells were seeded at 960,000 cells per well in 30×100mm Nunc dishes and differentiated as described above. The culture media was then replaced with 15 mL of DMEM:F12 complete medium supplemented with 0.1 % BDNF (6 dishes as untreated control) and 50 mM GBCA (15 dishes) or 50 mM BCA (9 dishes) for 48 h. Media was aspirated and cells were washed twice with 1× PBS. 200 μL of cold Triethylammonium bicarbonate (TEAB) was added to each plate and cells scraped off with a cell lifter. Cells were pooled (Control, 2× dishes; GBCA, 5× dishes; BCA, 3× dishes per replicate) and transferred to 1.5 mL tubes. 50-75 μL of each sample was collected in separate tubes for protein assay. To the remaining lysed cells, 1/10 (v/v) of 10 % sodium deoxycholate was added (1 % final concentration). Cells were snap frozen and stored at -70°C until TMT analyses.

All nine samples were processed for mass spectrometric analysis in a same batch using S-Traps and manufacturer’s protocol (Protifi, USA). Briefly, as the volume of the samples were large and variable across the samples, the samples were dried using vacuum centrifugation and then reconstituted in S-Trap Lysis buffer (50 mM TEAB, 5 % SDS), sonicated for 30s, boiled at 95 °C for 5 min and then cooled to room temperature.

An equal amount of each sample (100 μg) was aliquoted, and volumes normalised across all the samples using lysis buffer. Protein disulphide bridges were reduced by addition of DTT (final concentration 10 mM) at 60°C for 30 min, and then alkylated by addition of IAA (final concentration 25 mM) for 30 min in dark at ambient temperature. The pH of the samples was adjusted using aqueous phosphoric acid and diluted using S-Trap binding buffer (90 % aqueous methanol containing a final concentration of 100 mM TEAB, pH 7.55). The sample mixture was transferred to a labelled S-Trap column and centrifuged at 4,000× g, after which the flowthrough was discarded. The column was washed twice using S-Trap binding buffer and proteins retained on the column were digested in the presence of 125 μL trypsin solution (1:20 trypsin to protein ratio, total 5 μg total trypsin in 50 mM TEAB) for 1 h at 47°C.

Following digestion, peptides were eluted from the column after an addition of 50 mM TEAB and centrifugation. Remaining peptides were eluted from the column using a sequential centrifugation with addition of 0.2 % aqueous formic acid followed by 50 % aqueous acetonitrile containing 0.2 % formic acid. Peptides were dried by vacuum centrifugation and then reconstituted in 200 mM HEPES (pH 8.8). Peptide concentration was determined using the Pierce quantitative colorimetric peptide assay (Thermo Scientific, USA).

#### Sample labelling

Equal total peptide quantities from each of 10 (9 samples and a pool) sample were used for subsequent sample processing. A pooled sample was generated by mixing a small part of eight samples (GBCA Toxin Replicate 3 was not used due to limited amount of sample). Ten samples were labelled in a 10-plex TMT label batch. TMT reagent (Thermo Scientific, USA) labelling of each sample was performed using manufacturer’s protocol.

Briefly, anhydrous acetonitrile was added to each TMT label vial followed by vortexing and brief centrifugation. Aliquots of individual peptide samples were labelled with one of the individual TMT labels (total of ten labels) at room temperature for 1 h. To quench the excess TMT label in the sample, 5 % hydroxylamine was added to each sample, vortexed briefly and then incubated at room temperature for 15 min. Before pooling the samples, a ‘label check’ experiment was performed to ensure equal amounts of total peptide were pooled from all samples. The label check was performed by mixing small, equal aliquots of each individually labelled TMT sample and the mixed sample was vacuum dried using a vacuum centrifuge. Samples were reconstituted in 2 % acetonitrile, 0.1 % formic acid and analysed by LC-MS/MS. A normalization factor was obtained from the label check experiment and the original TMT-labelled peptide samples were then pooled at an equal, 1:1 ratio across all individual samples in the respective set.

#### Offline high pH reversed phase (HpH) fractionation of TMT labelled peptides

The pooled peptides were cleaned using a reverse-phase C18 clean-up column (Sep-pak, MA, USA) and dried in a vacuum centrifuge. The peptide mixture was resuspended in loading buffer (5 mM ammonia solution, pH 10.5), then separated into a total of 96 fractions using an Agilent 1260 HPLC system equipped with a fraction collector. Peptides were separated over a 55 min linear gradient from 3 to 30 % acetonitrile in 5 mM ammonia solution (pH 10.5) at a flow rate of 0.3 mL/min on a C18 reverse phase column (Agilent Technologies, Santa Clara, CA, USA). Finally, the 96 fractions were consolidated into 20 individual fractions, dried by vacuum centrifugation, then reconstituted in 0.1 % formic acid for LC-MS/MS analysis.

#### Data dependent acquisition (DDA) LC-MS/MS

HpH fractionated, TMT labelled peptides were subjected to LC-MS/MS analysis. Briefly, each peptide fraction was injected onto the peptide trap and washed with loading buffer for 10 min. The peptide trap was then switched in line with the analytical nano-LC column. Peptides were eluted from the trap onto the nano - LC column and peptides were separated with a linear gradient of 5 % mobile phase B to 30 % mobile over 110 min at a flow rate of 0.3 mL/min, followed by 85 % B for 8 min.

The column eluent was directed into the ionization source of the mass spectrometer operating in positive ion mode. Peptide precursors from 350 to 1850 *m/z* were scanned at 60 k resolution. The 10 most intense ions in the survey scan were fragmented by HCD using a normalized collision energy of 33 with a precursor isolation width of 0.8 *m/z*. Only precursors with charge state +2 to +5 were subjected to MS/MS analysis. The MS method had a minimum signal requirement value of 3000 ions for MS2 triggering, an AGC target value of 1×10^5^ for MS2 and a maximum injection time of 85 ms. MS/MS scan resolution was set at 45,000 and dynamic exclusion was set to 30s.

#### Protein identification and quantitation

The raw data files were processed using Proteome Discoverer (Version 2.1.0.81, ThermoFisher Scientific). Proteomics data were mapped and identified using SequestHT and Mascot against peptide sequence database for the *Homo sapiens* (database containing reviewed and non-reviewed proteins, downloaded from UniProt). The parameters for the data processing were as follows: Enzyme: Trypsin; Maximum missed cleavages: 2; Precursor mass tolerance: 20 ppm; Fragment mass tolerance: 0.02 Da; Dynamic modifications: Oxidation (M), Deamidated (N, Q), Glu->PyroGlu, Gln->Pyro-Glu, Acetyl (Protein N-Terminus), and TMT6plex (K) and TMT6plex (N-term); Static Modification: Carbamidomethyl (C); FDR and result display filters: Protein, Peptide and PSM FDR<1 %, Master proteins only. The processed quantitative data were used for the subsequent analysis.

### Differential expression analysis of proteomics data

Fold-change of proteins were determined using pairwise relative-abundance and comparison was performed across all the sample groups of results using *t-*Tests with the cut-off of log fold-change (log_2_FC) > 0.5 or < -0.5, and *p*-value < 0.05 for statistical significance. The UniProtIDs of proteins were then matched to the *H. sapiens* Ensembl annotations (EnsDb.Hsapiens.v86) to facilitate the identification of significant differential expressed proteins using in-house R scripts.

### Gene ontology (GO) and protein interaction network analysis of proteomics data

Gene ontology (GO) enrichment analysis were performed using the function *EnrichGO* in clusterProfiler v4.4.4 in R against *H. sapiens* Gene Ontology database *org.Hs.eg.db* acquired from Bioconductor [28,29]. The *p*-values were corrected using Benjamini-Hochberg (BH) method then the significance cut-off was set to adjusted *p*-value < 0.05. The interaction network nodes and edges of significant Molecular Function (MF) GO were generated using *cnetplotJam* function from the multienrichJam package (https://github.com/jmw86069/multienrichjam). The interactive networks were then visualized using Cytoscape R (Rcy3) v2.16.0 and igraph v1.3.2 (https://igraph.org/r/) [30].

### Cell cycle analysis

SH-SY5Y cells were plated on 60mm Nunc petri dishes (4×10^6^ cells in 5 mL DMEM, 10 % HI-FBS, 1 % penstrep per dish) and cultured for 24 h in a humidified, 5 % CO_2_, 37 °C incubator. The next day, the media was replaced with 3 mL fresh media supplemented with water, 25 mM BCA or 25 mM GBCA (three replicates per treatment) and the cells were incubated for 24 h, after which the cells were harvested and fixed in 1 mL of 70 % EtOH in 1× PBS at -20°C for at least 16 h. The EtOH was replaced with 1 mL of 1× PBS, supplemented with 0.1 μg/μL RNAse and 0.001 μg/μL of DAPI and the samples were incubated for 1 h at room temperature in dark. The samples were analysed by BD LSRFortessa^TM^ X-20 flow cytometry and FSC, SSC and V450/50 parameters for single cell population (10,000 events) for each sample were recorded using BD FACSDiva software. Cell cycle statistics were analysed using Watson’s model available in FlowJo software (version 10.8.1). Statistically significant difference (α = 0.01) among the treatments was analysed using Tukey’s HSD test available in SPSS software (version 28.0.1.0).

### Statistical analysis

All assays utilised three biological replicates and the values are represented as mean ± SEM. Cell confluence and viability data were statistically analysed using two-way ANOVA with multiple comparisons. All statistical analysis was performed using GraphPad Prism software 9.0 (GraphPad Software, Inc., La Jolla, USA). P < 0.05 was considered as statistically significant.

## Results

### BCA and GBCA dosage response of differentiated SH-SY5Y cells

Initially, we observed after 48 h of either 50 mM β-cyano-L-alanine (BCA) or γ-glutamyl-β-cyano-L-alanine (GBCA) treatment, differentiated SH-SY5Y cells appeared shrunken and had shorter dendrites, compared to the negative control (Figure 1a). The BCA treated cells were much smaller than the GBCA treated cells (Figure 1a). This indicated that BCA and GBCA might induce cell toxicity at different concentrations.

Next, we determined BCA or GBCA dosage responses of differentiated SH-SY5Y cells by treating the cells with a range of toxin concentrations for 48 h. SH-SY5Y cell differentiation was first confirmed by western blotting (Figure 1b) for Neurofilament H marker and cell proliferation was recorded by measuring cell confluence every 2 h during the treatment period. Cell viability was measured after 48 h using an Alamar Blue assay (Figure 1c, d). We observed an initial reduction in cell confluence after replacing the media with fresh (controls) or toxin supplemented media (time point 0). The cell confluence was stabilized after about 6 hours and only slightly increased throughout the treatment period in the controls (Figure 1c).

When treating the differentiated SH-SY5Y cells at BCA concentrations ranging from 20 to 40 mM, both cell confluence and cell viability were not significantly different to that of the control (Figure 1c, d). However, at 50 mM BCA the cells showed a significant (two-way ANOVA, p<0.01) reduction in cell confluence after 38 h of treatment (Figure 1c). At 60 mM BCA, cell confluence was significantly reduced from the start of the treatment and remained lower throughout the treatment period (Figure 1c). At 48 h, cell confluences of the 50 mM and 60 mM BCA treatments were significantly lower (51 ± 4.7 % and 42 ± 5.5 %, respectively) than the control (65 ± 6.3 %). As expected, cell viability analysis at 48 h using the Alamar Blue assay showed toxicity effects of BCA at 50 and 60 mM (Figure 1d). At 50 mM BCA, cell viability was reduced by about 50 % and at 60 mM BCA, cell viability was reduced by 92 % compared to the control (Figure 1d). In summary, a clear toxicity effect of BCA on differentiated SH-SY5Y cells was observed at 50 mM.

When treating the differentiated SH-SY5Y cells with 20 mM GBCA, both cell confluence and cell viability were not significantly different compared to the control after 48 h, despite a reduction in cell confluence for the first 6 h of the treated sample (Figure 1c, d). When the cells were treated with either 40, 60 or 80 mM GBCA, there was a significant reduction in the cell confluence at the start of the treatment and continued to be lower than the control throughout the treatment period (Figure 1c). After 48 hours, the cell confluences were 47 ± 5.1 %, 38 ± 5.0 % and 29 ± 2.7 % for the 40, 60 and 80 mM GBCA treatments, respectively, which were significantly (two-way ANOVA, p<0.01) lower compared to that of the control confluence of 60 ± 6.2 %. As expected, cell viability analysis at 48 h using the Alamar Blue assay showed toxicity effects of GBCA started at 40 mM (Figure 1d). Cell viability was reduced by about 30 % in the 40 mM GBCA treatment, and by about 60 and 90 % in the 60- and 80 mM treatments, respectively, compared to the control (Figure 1d). In summary, increasing GBCA concentration above 40 mM linearly decreased the cell confluence and cell viability of differentiated SH-SY5Y cells. We reviewed our BCA and GBCA cell confluence and viability response curves to identify the optimal concentrations for our TMT proteomics analysis. We decided to use 50 mM BCA and GBCA based on the LD_50_ in cells treated for 48 hours, as we also aimed to strike a balance between preserving cell survival while accurately assessing the impact of these compounds on cellular proteins.

### BCA and GBCA dosage response of primary mouse neurons

To test whether BCA or GBCA affects primary mouse embryonic neurons similarly to that of differentiated SH-SY5Y cells, we treated primary mouse neurons with similar BCA or GBCA concentrations and analysed the cell viability after 48h using Alamar Blue assay. Interestingly, treating the cells with BCA at concentration from 20 to 60 mM did not show any statistically significant reduction in cell viability compared to that of the control (Figure 2). However, we observed the GBCA toxicity effect on primary mouse neuron cells started as low as 20 mM (1.3 ± 0.46×10^4^ RFU) with a 75 % reduction in cell viability compared to that of the control (5.2 ± 0.07×10^4^ RFU) (Figure 2).We observed that primary mouse neurons are less sensitive to BCA toxin but more sensitive to GBCA toxin (toxicity effect started at 20 mM) when compared to differentiated SH-SY5Y cells, with the GBCA toxicity effect detectable at 40 mM (Figures 1 and 2).

**Figure 2.**
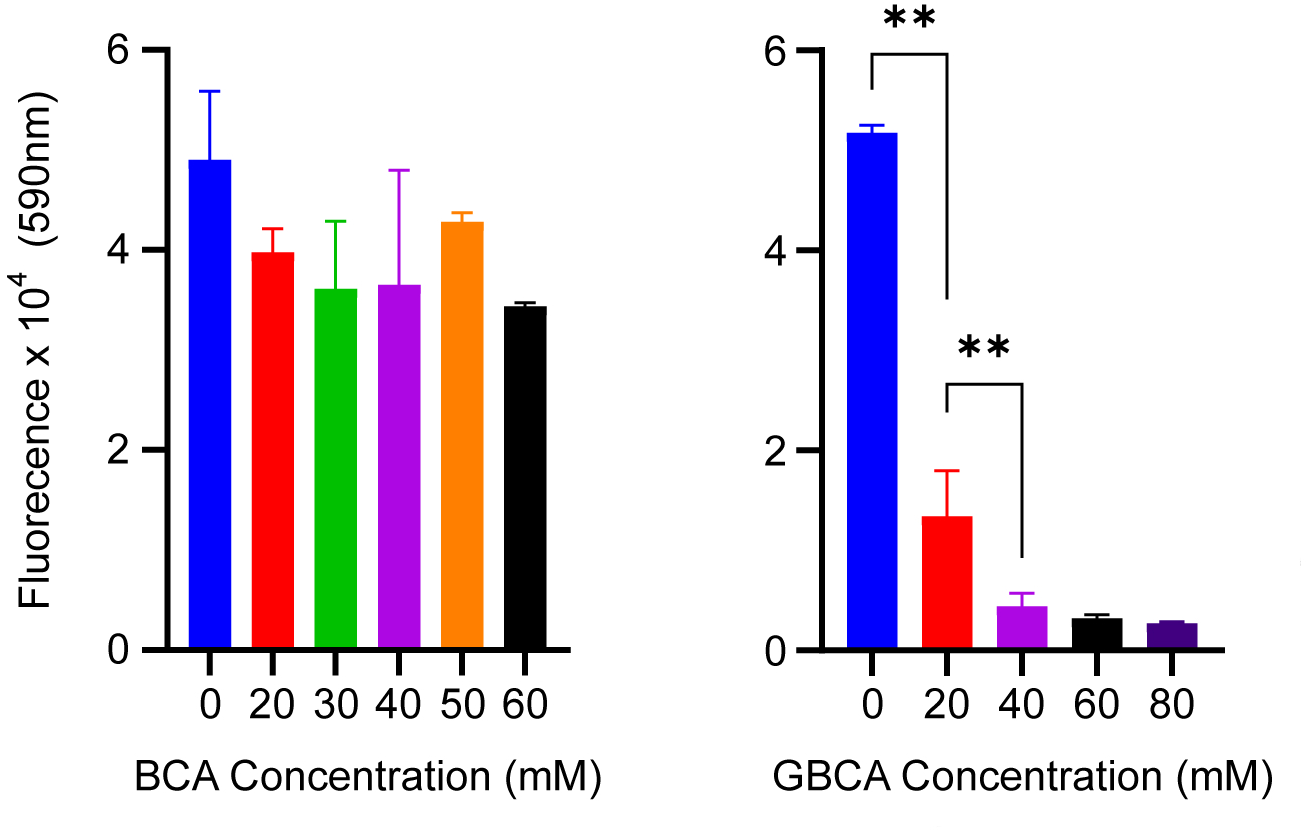
Cell viability of mouse primary neurons cultured in the absence or presence of BCA or GBCA for 48 h as determined by Alamar Blue assay. Data shown as mean ± SEM of triplicate biological replicates, ** *p*-value<0.01.

### Potential synergistic effects of BCA with GBCA

Both BCA and GBCA are present in *V. sativa* seeds and the compounds may act synergistically to induce toxicity within cells. First, we measured the BCA and GBCA concentrations within seed extracts of varieties Timok and Lov-2 and then calculated their molar ratios. Timok seeds contained 1 mM BCA and 4.2 mM GBCA and Lov-2 seeds 1.1 mM BCA and 4.2 mM GBCA as determined by LC-MS. The BCA and GBCA ratio in the seeds of both varieties was 1:4. In subsequent neural toxicity experiments using purified BCA and GBCA, we used a 1:4 BCA:GBCA ratio.

We tested the synergistic effect of different concentrations of purified BCA and GBCA (1:4 ratio) on differentiated SH-SY5Y cells. Cell confluence recorded every 6h for a total of 72h of treatment showed the highest concentration of 10 mM BCA/40 mM GBCA (-34±10.0 %, *p-value =* 0.0297) resulted in a significant decline in cell confluence 60 h of treatment compared to 10 mM BCA (13±4.42 %) or 40 mM GBCA alone (-21.6±3.71 %). Conversely lower BCA/GBCA concentration of 2.5 mM BCA/10 mM GBCA (20±10.9 %, *p-value* > 0.999) and 5 mM BCA/20 mM GBCA (-4.4±3.08 %, *p-*value > 0.999) showed no significant difference in cell confluence compared to the individual compounds alone (2.5 mM BCA (26.5±9.34 %), 5mM BCA (12.4±3.2 %), 10 mM GBCA (11±2.28 %), or 20 mM GBCA (2.5±3.53 %). No consistent synergistic relationship between BCA and GBCA at any concentration of the 1:4 ratio was observed, indicating an additive relationship between them. This suggests that any combined disruptive effects on cell confluence arise from the individual toxic effects of each compound rather than any additional interactions between them.

### Identification of differentially expressed proteins in BCA and GBCA treated differentiated SH-SY5Y cells

TMT quantitative proteomics was performed to identify differentially expressed proteins (DEPs) in differentiated SH-SY5Y cells treated with GBCA or BCA. A total of 6,827 proteins were quantified across the control, BCA and GBCA treated samples. For further analysis, a log_2_ fold change ratio of 0.5 and Wilcoxon test p < 0.05 was used. Amongst the 6,827 proteins, while 26 and 73 proteins (DEPs) were significantly up- and downregulated, respectively, in BCA-treated cells compared to the control (Figure 4a left, b left), 76 and 86 were significantly up- and down-regulated, respectively, in the GBCA-treated cells against the control (Figure 4a right, b right) (Table 1).

**Figure 3.**
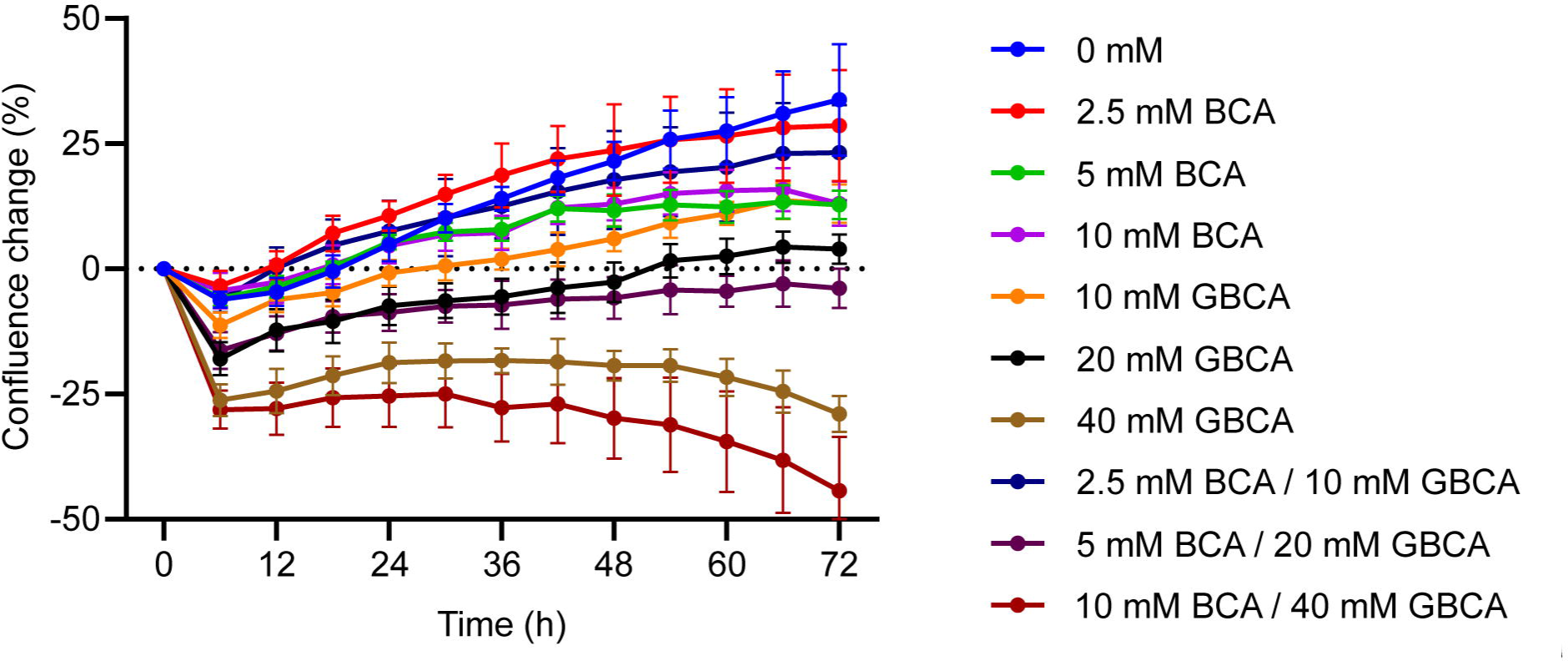
Real-time cell viability of differentiated SH-SY5Y cells cultured in the absence or presence of different concentrations of BCA, GBCA or combination of BCA and GBCA (1:4 BCA: GBCA). Cell viability was determined using an IncuCyte S3 system. Data shown as mean ± SEM of triplicate biological replicates.

**Figure 4.**
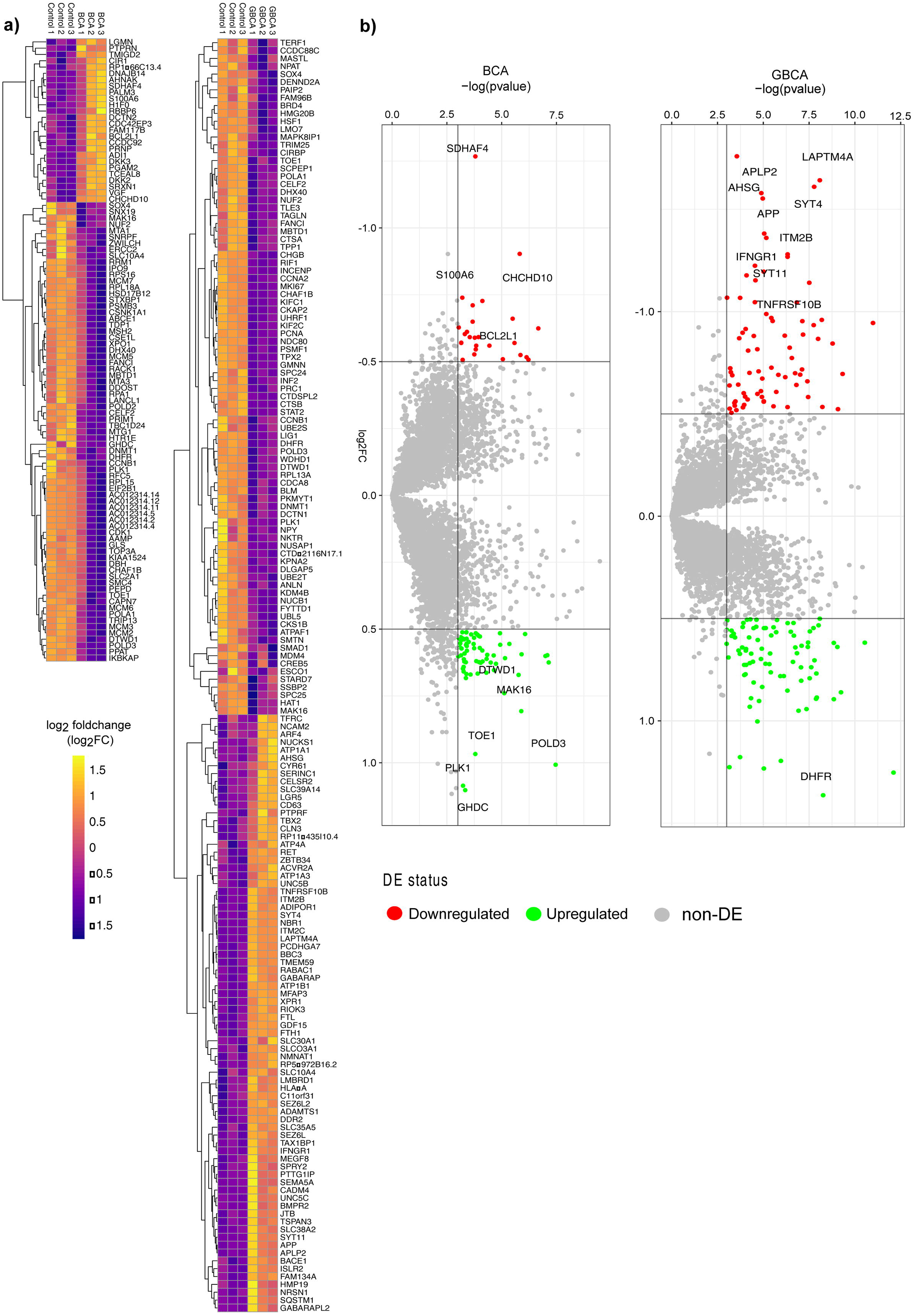
Protein expression profiling using a Tandem Mass Tagging quantitative proteomics approach. a) Unsupervised hierarchical clustering analysis of 99 differentially expressed proteins in differentiated SH-SY5Y cells treated with 50 mM BCA (left) and 162 differentially expressed proteins in cells treated with 50 mM GBCA (right) for 48 h. Colour key indicates the log_2_ expression level. b) Volcano plots of log_2_ fold change (x-axis) against log p-value (y-axis) of all 6,827 quantified proteins. Up- and downregulated proteins are coloured green and red, respectively.

### GO Enrichment analysis

Following identification of significantly dysregulated proteins, we performed enrichment analysis of gene ontology (GO) functional annotation terms utilizing the GO categories of cellular composition (CC), molecular function (MF), and biological process (BP). To detect significantly enriched biological functional categories, we performed GO enrichment analysis and ranked terms by enrichment score (− log_10_ adjusted *p* value < 0.05). Whereas 21 molecular function (MF), 31 cellular components (CC), and 103 biological processes (BP) GO terms for the differential expressed proteins were identified in BCA treated cells (Figure 5), 24 MF, 64 CC, and 194 BP GO terms were enriched in the differentially expressed proteins in GBCA treated cells. The top 10 significantly enriched MF, CC, and BP terms for BCA and GBCA treatments are shown (Figure 5a, b). DEPs showed starkly different enrichment across all GO categories between the BCA and GBCA treatments. In the BCA vs control, processes such as rRNA binding, DNA replication and other ribosome-related functional terms were enriched (Figure 5a). However, in the GBCA vs control, microtubule binding, tubulin binding and chromosome segregation processes were enriched (Figure 5b).

**Figure 5.**
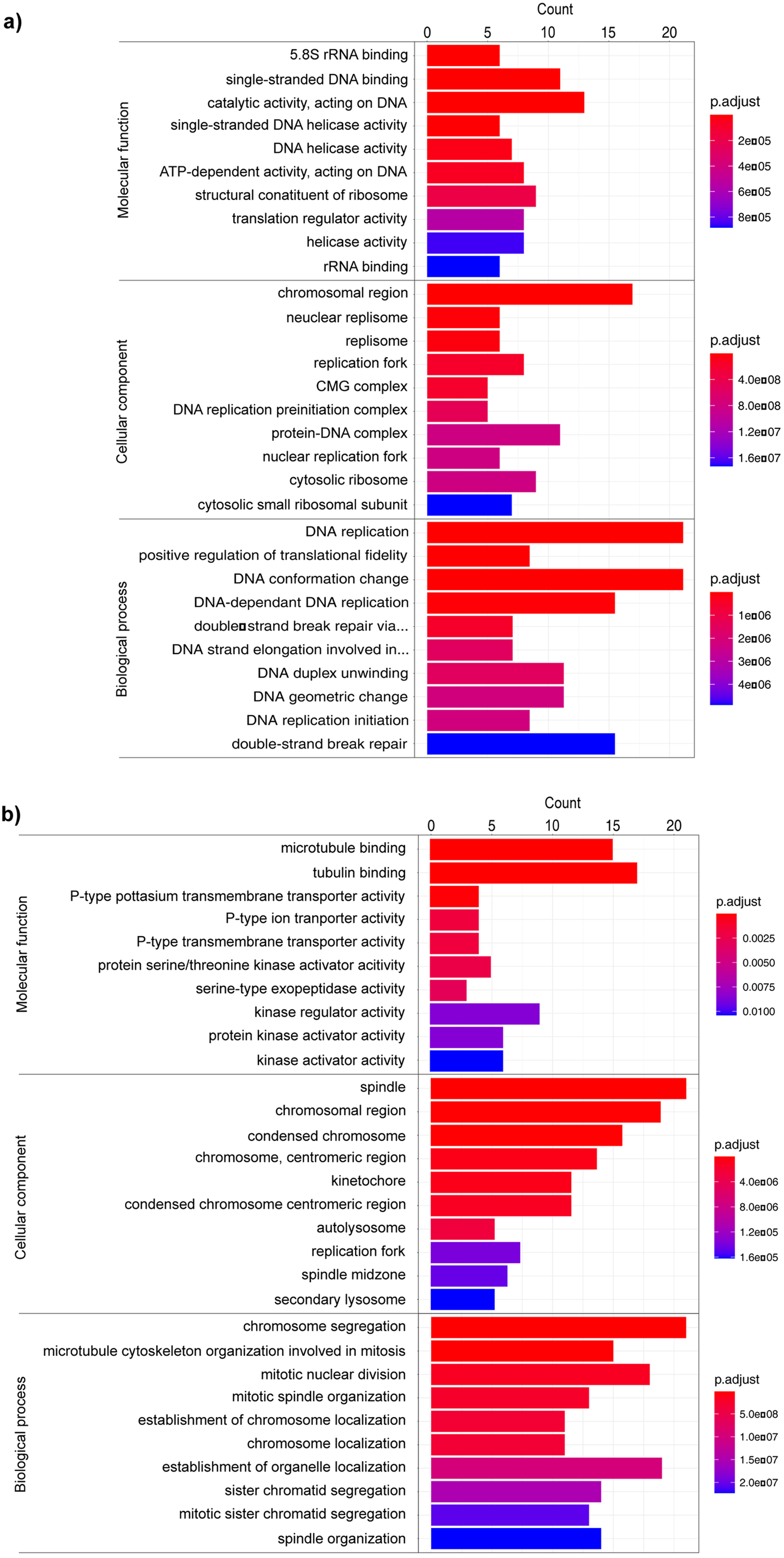
GO enrichment analysis for the top 10 significantly enriched molecular function (MF), cellular component (CC), and biological process (BP) terms for differentiated SH-SY5Y cells treated with a) 50 mM BCA or b) 50 mM GBCA. The y-axis denotes the categories of GO terms. The x-axis denotes the total protein count.

To better understand the differentially expressed proteins in either the BCA or GBCA vs control comparisons, we performed protein network analysis (see methods) to understand the implication of BCA and GBCA on molecular pathways (Figure 6). In the BCA vs control comparison, the top eight enriched molecular functions were derived from 20 proteins; interestingly all of which were upregulated. The key enriched networks were shown to be related to helicase activity, DNA binding and ribosome. For the GBCA vs control, 30 DEPs were linked to the top eight enriched molecular functions PPI networks including tubulin binding, kinase activity, P-type transporter activity and exopeptidase activity.

**Figure 6.**
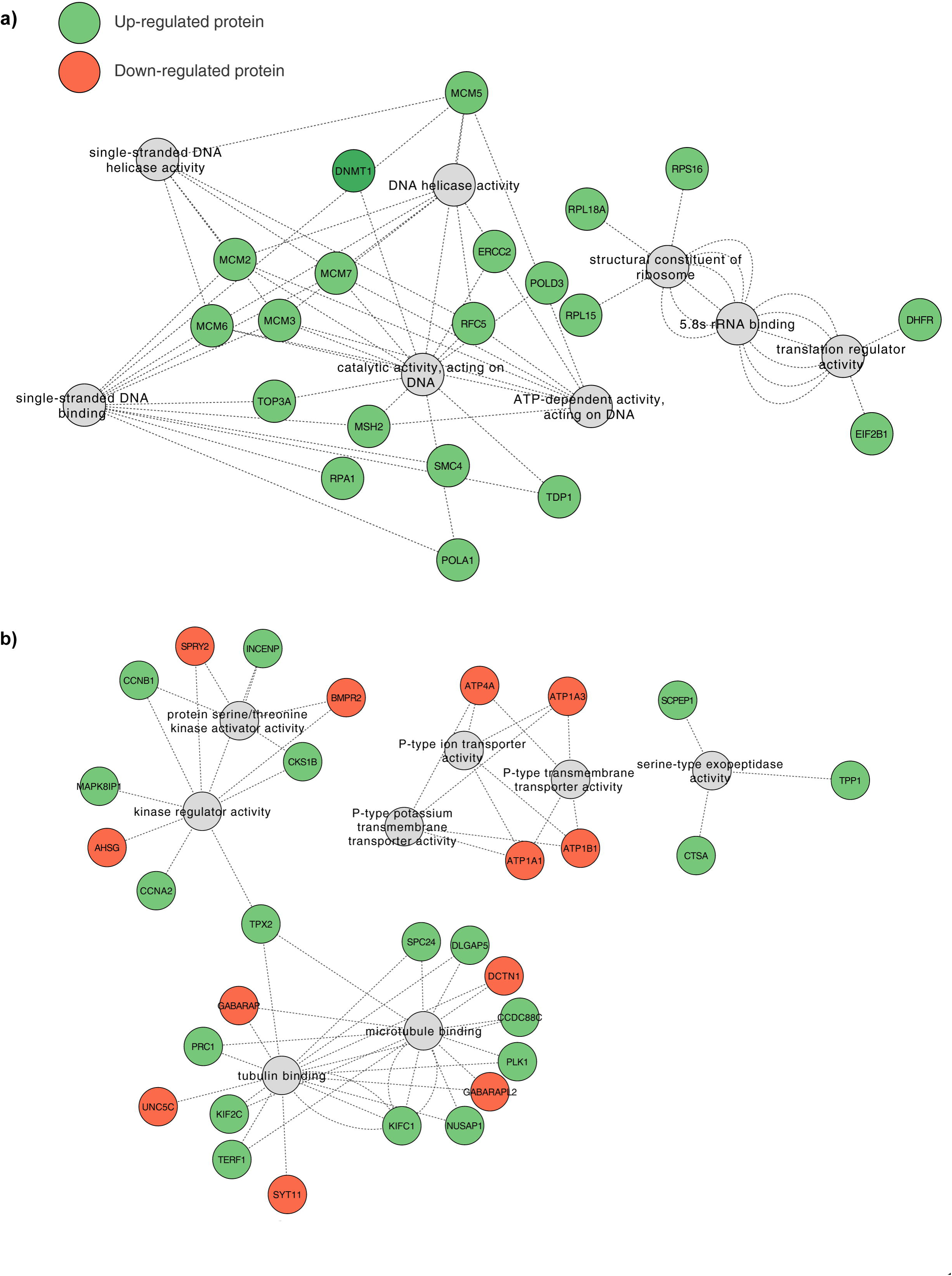
Predicted protein-protein interaction networks of differentially expressed proteins using Cytoscape. a) BCA/control. b) GBCA/control. Expression was assessed in differentiated SH-SY5Y cells treated with 50 mM BCA or 50 mM GBCA for 48 h.

### Effect of BCA and GBCA on SH-SY5Y cell cycle

GO enrichment analysis for DEPs in BCA or GBCA treated compared to untreated differentiated SH-SY5Y cells indicated that differential expression of proteins associated with different cellular processes may be contributing to the BCA- and GBCA-mediated cellular toxicity. We specifically observed enrichment of cell division related processes in the GBCA treated cells (Figure 5b). To validate this observation, we parsed undifferentiated SH-SY5Y cells treated with 25 mM GBCA for 24 h using flow cytometry to investigate changes in cell cycle pattern. We also analysed 25 mM BCA treated cells to determine if this had any covert effects on the cell division.

First, cell cycle analysis showed no significant change in G_0_-G_1_ cell population (65.03±1.88 %) in BCA treated than the untreated cells (61.87±1.99 %). However, we observed a small but non-significant increase in number of cells in the S phase of BCA treated (31.20±0.46 %) than untreated (26.63±0.63 %) cells, and a reduced number of cells that entered G_2_-M phase (5.09±0.41 %) in BCA treated than the control (26.63±0.63 %) cells (Figure 7a, b). Second, the cell cycle analysis suggested that GBCA treatment significantly altered the cell cycle profile. We observed a significant increase in G_0_-G_1_ cells in GBCA treated (77.63±0.32 %) compared to the untreated cells (61.67±1.99 %). In addition, we also observed a decrease in the number of cells in S (23.53±0.20 %) and G_2_-M phase (4.49±0.29 %) in GBCA treated than the control cells (26.63±0.63 % and 8.53±0.56 %, respectively) (Figure 7a, b).

**Figure 7.**
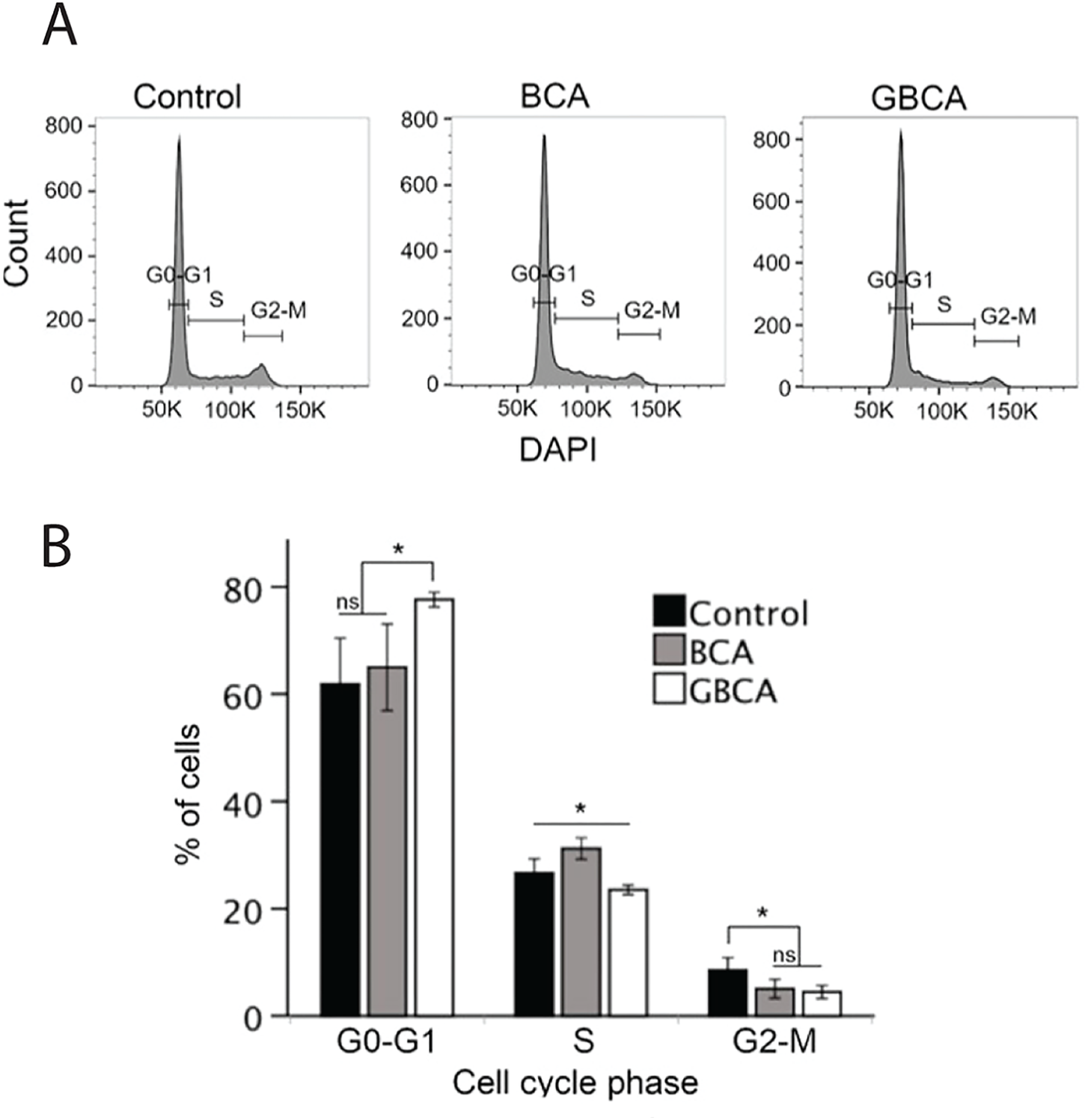
SH-SY5Y cells displayed cell cycle shift after 24 h treatment with BCA or GBCA. a) Cell morphology of control, BCA (25 mM) or GBCA (25 mM) treated cells. Scale bar = 200 µm. b) Cell cycle analysis of control, BCA or GBCA treated cells using flow cytometry (a representative result from three replicates). c) Percentage of cells at different phases of cell cycle of control, BCA or GBCA treated cells. Data shown as mean ± SEM of three biological replicates, \**p*-value<0.01, ns = not significant.

### Utilisation of *V. sativa* seed extracts for neurotoxicity assessment

To assess the capability of the *in vitro* SH-SY5Y-based assay on more complex matrices, filtered and unfiltered extracts of *V. sativa* cultivars Lov-2 and Timok were applied to differentiated cells for 72 h (Figure 8b), and both were compared to the pure GBCA standards (Figure 8a). To ensure that the concentrations of GBCA standards used in the cell viability assay fall within the range of all seed extract samples, we analysed *V. sativa* crude extracts using HPLC-MS, which determined GBCA concentrations of 4.17 mM for Lov-2 and 4.34 mM for Timok cultivars, respectively. These values fall within the range of the purified GBCA standard curve utilised in the cell viability assay. Filtration (3 kDa) of both Lov-2 (5.2±0.56×10^4^ nm, *p* = 0.0211) and Timok (6.5±0.727×10^4^ nm, *p* ≤ 0.0002) extracts prior to treatment caused a significant increase in cell viability compared to the unfiltered Lov-2 (2.9±0.389×10^4^ nm) and Timok (2.1±0.245×10^4^ nm) extracts. Furthermore, a comparison was undertaken between the cell viability of filtered *V. sativa* extracts and the cell viability standard curve established from GBCA-treated cells (y = -9.994ln(x) + 115.82, R² = 0.8356). This analysis provided the average estimated GBCA concentrations in Lov-2 (7.24±1.1 mM) and Timok (5.15±1.10 mM) seed extracts (Figure 8a, b), consistent with those determined through HPLC-MS quantification.

**Figure 8.**
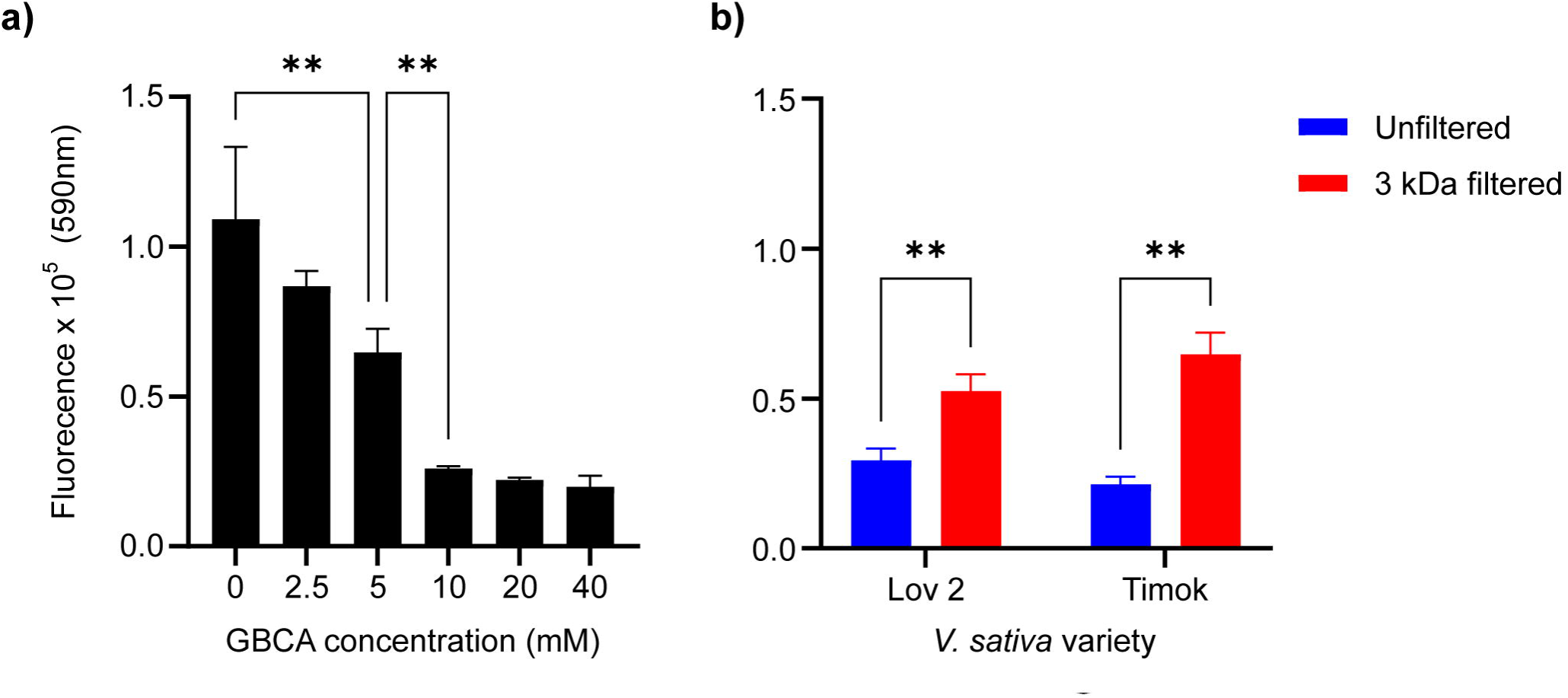
Relative cell viability of differentiated SH-SY5Y cells treated with a) varying concentrations of GBCA or b) unfiltered or 3 kDa filtered seed extracts from either ground V*. sativa* var. Lov-2 or Timok. Relative cell viability was determined at 72 h after the treatment using Alamar Blue assay (emitting fluorescence from live cells were measured at 590 nm). GBCA concentration in Lov-2 and Timok seed extracts were estimated 4.17 mM and 4.34 mM, respectively, using LC-MS. Data shown as mean ± SEM of three biological replicates, \*\**p*-value<0.01.

## Discussion

BCA and GBCA are the two most potent antinutritional compounds present in the seeds of the underutilized legume *V. sativa*. Animals have been shown to exhibit typical neurotoxicity symptoms when fed on *V. sativa* seed [31] supplemented diet or intravenous application of pure BCA or GBCA [11,32]. Evidence for the metabolic mechanisms underlying these neurotoxicity symptoms is sparse. To investigate the mechanisms, we first developed a BCA and GBCA *in vitro* neurotoxicity assay using retinoic acid differentiated SH-SY5Y cells and then investigated the potential mechanism of how BCA and GBCA cause toxicity by assaying the effect of these toxins on the proteome of SH-SY5Y-derived neurons using TMT-based quantitative proteomics approach. Our findings suggested many significant DEPs related to DNA helicase, DNA binding activity and the ribosome were impacted in BCA treated cells, whilst a significant DEPs related to tubulin binding, kinase activity, P-type transporter activity and exopeptidase activity were affected in the GBCA treated cells. The dysregulation of these critical cellular pathways may indicate mechanisms by which BCA or GBCA individually or together that lead to previously reported neurotoxicity.

BCA is a cyanide containing, non-protein amino acid from which toxicity is thought not to stem from energy depletion by the release of free cyanide [15], but instead cystathionine γ-lyase (CSE) enzyme inhibition causing accumulation of free radicals and subsequent apoptosis by oxidative stress is accepted as the most likely mechanism [12]. Oxidative stress is commonly associated with DNA damage phenotypes [33,34] and in the current study, many DEPs associated with DNA damage were upregulated in BCA treated SH-SY5Y-derived neurons. The minichromosome maintenance (MCM) family (MCM2,3,5,6,7) of proteins involved in DNA helicase activity [35] were all significantly upregulated. BCA-mediated oxidative stress can cause genomic instability through the action of cyclin-dependent kinases (CDKs) [36] as observed in many cancers [37,38]. Single stranded DNA binding proteins were significantly upregulated such as TDP1 (responsible for DNA repair following oxidative damage) [39], MSH2 (engaged in DNA mismatch repair) [40], SMC4 (responsible for replication damage repair) [41] and RFC5, (necessary for slowing of DNA synthesis when DNA is damaged during replication) [42]. The high proportion of DEPs associated with DNA repair could indicate a failed cell cycle checkpoint response (G_2_/M checkpoint), the effects of these DEPs were further supported by flow cytometry results assessing the cell cycle distribution of BCA treated undifferentiated SH-SY5Y cells. In cells treated with BCA, the cell cycle was significantly impeded at the transition step from S phase to G_2_-M phase, therefore increasing the number of S phase cell population. The proportion of cells experiencing S phase cell cycle arrest [43] could initiate apoptosis of cells [44] in the event of unsuccessful DNA repair during the arrest. This process could be occurring in the BCA treated SH-SY5Y cells but may not be the sole contributor to neurotoxicity based on our observation as the increased proportion of S phase cells was not significant (*p-value* > 0.05).

We also observed upregulation of several structural ribosomal proteins, including ribosomal protein large subunits (RPL) RPL15 and RPL18A and ribosomal protein small subunit (RPS) RPS16 in BCA treated cells. That these cellular processes are known to be responsible for successful transcription, translation, and subsequent proliferation of cells [45], are upregulated during periods of oxidative stress and are essential for maintaining ribosomal structure and protein synthesis [46] is consistent with our GO analysis of BCA DEPs, and suggestes that the presence of BCA in SH-SY5Y-derived cells enhances positive regulations of translation fidelity (Figure 5).

In contrast to BCA, GBCA seems to disturb more broadly the cell cycle regulation of differentiated SH-SY5Y cells. Even though we observed strong Neurofilament H differentiation marker expression in the differentiated SH-SY5Y cells (Figure 1b), presence of a small proportion of undifferentiated dividing cells could be the reason we identified a set of cell cycle related proteins in the GBCA treated SH-SY5Y cells. Many DEPs in GBCA treated cells were related to the maintenance of mitosis and the cell cycle as suggested by the GO and network analysis, which we then confirmed this with our flow cytometry results (Figure 7). The increase in G_0_-G_1_ cell population after GBCA treatment suggests that both the cell transition between G_0_-G_1_ phase to S phase and the G_2_-M phase are impeded, and this observation aligns with the GO enrichment terms identified in the DEPs analysis, which include microtubule binding, spindle organization, condensed chromosome, chromosome segregation, and microtubule cytoskeleton organization involved in mitosis. Specifically, nine proteins were found to be significantly upregulated, while five were significantly downregulated compared to the control (Figure 4b). Our network analysis suggested that several of these pathways cross-interact with one another (Figure 6). The observed dysregulation would likely initiate mitotic arrest and partial activation of apoptosis via p53 induction [47].

Interestingly, we observed that differentiated SH-SY5Y cells were sensitive to both BCA and GBCA, however the mouse primary embryonic neurons were mostly sensitive to GBCA. The reason for this difference is not clear, although the cytotoxicity and neurotoxicity of compounds can vary significantly between tested cell types [53,54]. That differences in the effect of BCA and GBCA could be due to the variation in physiological state of *in vitro* differentiated SH-SY5Y human cells (a proportion of which may still be cycling) vs highly homogeneous postmitotic mouse neurons cannot be ruled out. Together, these differences in BCA and GBCA sensitivity across differing cell types should be considered when planning future animal trials for assessment of *V. sativa* toxicity. However, we reason that due to the lower prevalence of BCA (1 mM) compared to GBCA (4.5 mM) in the raw vetch seeds and lack of observed synergistic toxicity effects between BCA and GBCA at those same ratios upon SH-SY5Y-derived cells, it is unlikely that this would affect future testing towards *V. sativa* as an animal food source.

### Conclusions

In this study, we used TMT-based quantitative proteomics to establish the underlying mechanisms of the effect of BCA and GBCA on differentiated SH-SY5Y cells. A total of 96 and 162 significantly DEPs were identified in BCA and GBCA treated differentiated SH-SY5Y cells, respectively, from the total 6,827 proteins. The GO enrichment analysis showed that BCA treatment resulted in significantly DEPs related to helicase activity, DNA binding and ribosome whilst GBCA treatment led to significantly DEPs related to tubulin binding, kinase activity, P-type transporter activity and exopeptidase activity. Upregulation of several MCM family and single stranded DNA binding proteins TDP1, MSH2, SMC4 and RFC5 by BCA treatment indicates potentially high levels of oxidative stress consistent with previously reported CSE inhibition and downstream accumulation of reactive oxygen species triggering apoptosis. Whilst additional experiments will provide further insights into the GBCA toxicity mechanisms, the observation of DEPs associated with previously described excitotoxicity pathways and neurotoxicity symptoms is encouraging. Our findings provide new insights relating the mechanistic action of plant-derived toxins from *V. sativa* seeds; BCA and GBCA in mammalian cell types. These results shed light into the putative mechanism of neurotoxicity caused by *V. sativa* toxins, explaining the observed neuronal cell death in monogastric animals when the toxins are present. Several DEPs identified have the potential to be useful biomarkers for future animal feed screening trials using zero toxin *V. sativa*.

## Acknowledgements/Funding, people

This work was supported by an Australian Research Council (ARC) grant LP200200957 awarded to IS, JG and RS, a South Australian Grains and Industry Trust (SAGIT) grant UA720 awarded to IS, a Biotechnology and Biological Sciences Research Council (BBSRC) grant BB/V018108/1 awarded to IF and Research Training Program scholarships (RTPS) from the University of Adelaide awarded to SR, RB, and PN.

## References

[1] L. Roberts, 9 Billion?, Science (1979). 333 (2011) 540–543. 10.1126/science.333.6042.540.

[2] FAO, The future of food and agriculture, Alternative pathways to 2050, 2018.

[3] P. Alexander, M.D.A. Rounsevell, C. Dislich, J.R. Dodson, K. Engström, D. Moran, Drivers for global agricultural land use change: The nexus of diet, population, yield and bioenergy, Global Environmental Change. 35 (2015) 138–147. 10.1016/j.gloenvcha.2015.08.011.

[4] K.A. Johnson, D.E. Johnson, Methane emissions from cattle, J Anim Sci. 73 (1995) 2483–2492. 10.2527/1995.7382483x.

[5] V. Nguyen, S. Riley, S. Nagel, I. Fisk, I.R. Searle, Common Vetch: A Drought Tolerant, High Protein Neglected Leguminous Crop With Potential as a Sustainable Food Source, Front Plant Sci. 11 (2020) 1–7. 10.3389/fpls.2020.00818.

[6] FAO, 11 Most Important Pulses FAO, (1994). http://www.fao.org/es/faodef/fdef04e.htm.

[7] J.T. Enopala, F.J. González, E. de La Barrera, Physiological responses of the green manure, vicia sativa, to drought, Bot Sci. 90 (2012) 305–311. 10.17129/botsci.392.

[8] S.C. Valentine, B.D. Bartsch, Production and composition of milk by dairy cows fed common vetch or lupin grain as protein supplements to a silage and pasture-based diet in early lactation, Aust J Exp Agric. 36 (1996) 633–636. 10.1071/EA9960633.

[9] Z. Mao, H. Fu, Z. Nan, C. Wan, Fatty acid, amino acid, and mineral composition of four common vetch seeds on Qinghai-Tibetan plateau, Food Chem. 171 (2015) 13–18. 10.1016/j.foodchem.2014.08.090.

[10] M.E. Tate, J. Rathjen, I. Delaere, D. Enneking, Covert trade in toxic vetch continues, Nature. 400 (1999) 207. 10.1038/22198.

[11] C. Ressler, Isolation and Identification from Common Vetch the Neurotoxin a Possible Factor in Neurolathyrism, J Biol Chem. 237 (1962) 733–736.

[12] M. Pfeffer, C. Ressler, β-Cyanoalanine, an inhibitor of rat liver cystathionase, Biochem Pharmacol. 16 (1967) 2299–2308. 10.1016/0006-2952(67)90217-1.

[13] C. Ressler, J.G. Tatake, E. Kaizer, D.H. Putnam, Neurotoxins in a Vetch Food: Stability to Cooking and Removal of γ-Glutamyl-β-cyanoalanine and β-Cyanoalanine and Acute Toxicity from Common Vetch ( *Vicia sativa* L.) Legumes, J Agric Food Chem. 45 (1997) 189–194. 10.1021/jf9603745.

[14] J.M. Rathjen, The potential for Vicia sativa L. as a grain legume for South Australia, University of Adelaide, 1997.

[15] D.N. Roy, M.I. Sabri, R.J. Kayton, P.S. Spencer, β-cyano-L-alanine toxicity: Evidence for the involvement of an excitotoxic mechanism, Nat Toxins. 4 (1996) 247–253. 10.1002/(sici)(1996)4:6<247::aid-nt1>3.0.co;2-m.

[16] D. Enneking, The toxicity of Vicia Species and their utilisation as grain legumes, University of Adelaide, 1994.

[17] F. Lambein, Y. Kuo, F. Ikegami, K. Kusama-eguchi, Grain legumes and human health, Genetics. 1 (2008) 422–432.

[18] M.E. Tate, D. Enneking, A mess of Red Pottage, Nature. 359 (1992) 357–358. 10.1038/22198.

[19] N. Ghasemi, H. Secen, H. Yılmaz, B. Binici, A.C. Goren, Determination of neurotoxic agents as markers of common vetch adulteration in lentil by LC-MS/MS, Food Chem. 221 (2017) 2005–2009. 10.1016/j.foodchem.2016.11.079.

[20] P. Thavarajah, D. Thavarajah, G.A.S. Premakumara, A. Vandenberg, Detection of Common Vetch (Vicia sativa L.) in Lentil (Lens culinaris L.) using unique chemical fingerprint markers, Food Chem. 135 (2012) 2203–2206. 10.1016/j.foodchem.2012.06.124.

[21] Y.F. Huang, X.L. Gao, Z.B. Nan, Z.X. Zhang, Potential value of the common vetch (Vicia sativa L.) as an animal feedstuff: a review, J Anim Physiol Anim Nutr (Berl). 101 (2017) 807–823. 10.1111/jpn.12617.

[22] J.E. May, J. Xu, H.R. Morse, N.D. Avent, C. Donaldson, Toxicity testing: the search for an *in vitro* alternative to animal testing, Br J Biomed Sci. 66 (2009) 160–165. 10.1080/09674845.2009.11730265.

[23] W.F. An, N. Tolliday, Cell-Based Assays for High-Throughput Screening, Mol Biotechnol. 45 (2010) 180–186. 10.1007/s12033-010-9251-z.

[24] B. Tschiersch, Occurrence of γ-glutamyl-β-cyanoalanine, Tetrahedron Lett. 5 (1964) 747–749. 10.1016/S0040-4039(00)90386-1.

[25] S. Kaech, G. Banker, Culturing hippocampal neurons, Nat Protoc. 1 (2006) 2406–2415. 10.1038/NPROT.2006.356.

[26] M.M. Shipley, C.A. Mangold, M.L. Szpara, Differentiation of the SH-SY5Y Human Neuroblastoma Cell Line, J Vis Exp. 2016 (2016) 53193. 10.3791/53193.

[27] C. Megías, I. Cortés-Giraldo, J. Girón-Calle, J. Vioque, M. Alaiz, Determination of β -Cyano-L-alanine, γ -Glutamyl-β -cyano-L-alanine, and Common Free Amino Acids in *Vicia sativa* (Fabaceae) Seeds by Reversed-Phase High-Performance Liquid Chromatography, J Anal Methods Chem. 2014 (2014) 1–5. 10.1155/2014/409089.

[28] G. Yu, L.-G. Wang, Y. Han, Q.-Y. He, clusterProfiler: an R Package for Comparing Biological Themes Among Gene Clusters, OMICS. 16 (2012) 284–287. 10.1089/omi.2011.0118.

[29] T. Wu, E. Hu, S. Xu, M. Chen, P. Guo, Z. Dai, T. Feng, L. Zhou, W. Tang, L. Zhan, X. Fu, S. Liu, X. Bo, G. Yu, clusterProfiler 4.0: A universal enrichment tool for interpreting omics data, The Innovation. 2 (2021) 100141. 10.1016/j.xinn.2021.100141.

[30] J.A. Gustavsen, S. Pai, R. Isserlin, B. Demchak, A.R. Pico, RCy3: Network biology using Cytoscape from within R, F1000Res. 8 (2019) 1774. 10.12688/f1000research.20887.1.

[31] C. Ressler, J.G. Tatake, E. Kaizer, D.H. Putnam, Neurotoxins in a Vetch Food: Stability to Cooking and Removal of γ-Glutamyl-β-cyanoalanine and β-Cyanoalanine and Acute Toxicity from Common Vetch ( Vicia sativa L.) Legumes, J Agric Food Chem. 45 (2002) 189–194. 10.1021/jf9603745.

[32] C. Ressler, J. Nelson, M. Pfeffer, Metabolism of beta-cyanoalanine, Biochem Pharmacol. 16 (1967) 2309–2319. http://www.ncbi.nlm.nih.gov.offcampus.lib.washington.edu/pubmed/6075393 http://www.sciencedirect.com.offcampus.lib.washington.edu/science?_ob=MImg&_imagekey=B6T4P-478FGV8-6N-1&_cdi=4980&_user=582538&_pii=0006295267902183&_origin=browse&_zone=rslt_list.

[33] A. Barzilai, K.-I. Yamamoto, DNA damage responses to oxidative stress, DNA Repair (Amst). 3 (2004) 1109–1115. 10.1016/j.dnarep.2004.03.002.

[34] K. Davies, The Broad Spectrum of Responses to Oxidants in Proliferating Cells: A New Paradigm for Oxidative Stress, IUBMB Life. 48 (1999) 41–47. 10.1080/713803463.

[35] K. Labib, J.A. Tercero, J.F.X. Diffley, Uninterrupted MCM2-7 Function Required for DNA Replication Fork Progression, Science (1979). 288 (2000) 1643–1647. 10.1126/science.288.5471.1643.

[36] S. Tanaka, Y.-S. Tak, H. Araki, The role of CDK in the initiation step of DNA replication in eukaryotes, Cell Div. 2 (2007) 16. 10.1186/1747-1028-2-16.

[37] J. Zhou, M. Wang, Z. Zhou, W. Wang, J. Duan, G. Wu, Expression and Prognostic Value of MCM Family Genes in Osteosarcoma, Front Mol Biosci. 8 (2021). 10.3389/fmolb.2021.668402.

[38] M. Das, S. Singh, S. Pradhan, G. Narayan, MCM Paradox: Abundance of Eukaryotic Replicative Helicases and Genomic Integrity, Mol Biol Int. 2014 (2014) 1–11. 10.1155/2014/574850.

[39] Y. Pommier, S.N. Huang, R. Gao, B.B. Das, J. Murai, C. Marchand, Tyrosyl-DNA-phosphodiesterases (TDP1 and TDP2), DNA Repair (Amst). 19 (2014) 114–129. 10.1016/j.dnarep.2014.03.020.

[40] M.A. Edelbrock, S. Kaliyaperumal, K.J. Williams, Structural, molecular and cellular functions of MSH2 and MSH6 during DNA mismatch repair, damage signaling and other noncanonical activities, Mutation Research/Fundamental and Molecular Mechanisms of Mutagenesis. 743–744 (2013) 53–66. 10.1016/j.mrfmmm.2012.12.008.

[41] Y. Wang, Z. Wu, The Clinical Significance and Transcription Regulation of a DNA Damage Repair Gene, SMC4, in Low-Grade Glioma via Integrated Bioinformatic Analysis, Front Oncol. 11 (2021). 10.3389/fonc.2021.761693.

[42] K. Sugimoto, S. Ando, T. Shimomura, K. Matsumoto, Rfc5, a replication factor C component, is required for regulation of Rad53 protein kinase in the yeast checkpoint pathway, Mol Cell Biol. 17 (1997) 5905–5914. 10.1128/MCB.17.10.5905.

[43] Z. Xu, F. Zhang, C. Bai, C. Yao, H. Zhong, C. Zou, X. Chen, Sophoridine induces apoptosis and S phase arrest via ROS-dependent JNK and ERK activation in human pancreatic cancer cells, Journal of Experimental and Clinical Cancer Research. 36 (2017). 10.1186/s13046-017-0590-5.

[44] I.I. Kruman, Why do Neurons Enter the Cell Cycle?, Cell Cycle. 3 (2004) 767–771. 10.4161/cc.3.6.901.

[45] D.A. Smagin, I.L. Kovalenko, A.G. Galyamina, A.O. Bragin, Y.L. Orlov, N.N. Kudryavtseva, Dysfunction in Ribosomal Gene Expression in the Hypothalamus and Hippocampus following Chronic Social Defeat Stress in Male Mice as Revealed by RNA-Seq, Neural Plast. 2016 (2016) 1–6. 10.1155/2016/3289187.

[46] A. Saha, S. Das, M. Moin, M. Dutta, A. Bakshi, M.S. Madhav, P.B. Kirti, Genome-Wide Identification and Comprehensive Expression Profiling of Ribosomal Protein Small Subunit (RPS) Genes and their Comparative Analysis with the Large Subunit (RPL) Genes in Rice, Front Plant Sci. 8 (2017) 1553. 10.3389/fpls.2017.01553.

[47] J.D. Orth, A. Loewer, G. Lahav, T.J. Mitchison, Prolonged mitotic arrest triggers partial activation of apoptosis, resulting in DNA damage and p53 induction, Mol Biol Cell. 23 (2012) 567–576. 10.1091/mbc.e11-09-0781.

[48] CC Walton Enríquez, Mitotic biology of primary neurons, Universidad Autónoma de Madrid, 2018.

[49] C.C. Walton, W. Zhang, I. Patiño-Parrado, E. Barrio-Alonso, J.-J. Garrido, J.M. Frade, Primary neurons can enter M-phase, Sci Rep. 9 (2019) 4594. 10.1038/s41598-019-40462-4.

[50] H. Wang, R.W. Olsen, Binding of the GABAA Receptor-Associated Protein (GABARAP) to Microtubules and Microfilaments Suggests Involvement of the Cytoskeleton in GABARAPGABAA Receptor Interaction, J Neurochem. 75 (2002) 644–655. 10.1046/j.1471-4159.2000.0750644.x.

[51] M. Ankarcrona, J.M. Dypbukt, E. Bonfoco, B. Zhivotovsky, S. Orrenius, S.A. Lipton, P. Nicotera, Glutamate-induced neuronal death: A succession of necrosis or apoptosis depending on mitochondrial function, Neuron. 15 (1995) 961–973. 10.1016/0896-6273(95)90186-8.

[52] D.A. McCormick, GABA as an inhibitory neurotransmitter in human cerebral cortex, J Neurophysiol. 62 (1989) 1018–1027. 10.1152/jn.1989.62.5.1018.

[53] D. Popova, J. Karlsson, S.O.P. Jacobsson, Comparison of neurons derived from mouse P19, rat PC12 and human SH-SY5Y cells in the assessment of chemical- and toxin-induced neurotoxicity, BMC Pharmacol Toxicol. 18 (2017) 42. 10.1186/s40360-017-0151-8.

[54] H.J. Heusinkveld, R.H.S. Westerink, Comparison of different in vitro cell models for the assessment of pesticide-induced dopaminergic neurotoxicity, Toxicology in Vitro. 45 (2017) 81–88. 10.1016/j.tiv.2017.07.030.

